# Presence or absence of Ras-dimerization shows distinct kinetic signature in Ras-Raf interaction

**DOI:** 10.1101/810150

**Authors:** Sumantra Sarkar, Angel E. Garcia

## Abstract

In eukaryotes, MAPK pathways play a central role in making several important cellular decisions, including cell proliferation and development of an organism. Ras, a small GTPase, interacts with the protein Raf to create activated Ras-Raf complex (Raf dimer) that activates the downstream effectors in the ERK pathway, one of the many MAPK pathways. Malfunctioning Ras-Raf “switches” cause almost 30% of all known cancer. Hence, understanding Ras-Raf interaction is of paramount importance. Despite decades of research, the detailed mechanism of Ras-Raf interaction is still unclear. It has been hypothesized that Ras dimerization is necessary to create the activated Raf dimer. Although there are circumstantial evidences supporting the Ras dimerization hypothesis, direct proof of Ras dimerization is still inconclusive. In the absence of conclusive direct experimental proof, this hypothesis can only be examined through indirect evidences of Ras dimerization. In this paper, using a multi-scale simulation technique, we provide multiple criteria that distinguishes an activation mechanism involving Ras dimerization from another mechanism that does not involve Ras dimerization. The provided criteria will be useful in the investigation of not only Ras-Raf interaction but also other two-protein interactions.

## INTRODUCTION

Ever since its discovery as an oncogene, Ras has been subject of intense scientific research^1–9^. Part of that effort identified the interaction of its protein product Ras with other proteins, in particular Raf, and contributed to the discovery of the cell signaling networks. Since then, Ras-Raf interaction has taken a center stage in cancer research and it has been established as a model system to study how mutated protein-protein interactions lead to cancer. In fact, mutated Ras-Raf interaction has been implicated in almost 30% of all known cancers and various other diseases^1^. Despite its notoriety, no universally effective drugs have been developed against Ras, leading the community to speculate that Ras is *undruggable*. However, this assessment reflects the frustration and the desperation of the scientific community, rather than being the scientific truth. Our inability to produce effective drugs stemmed from an incomplete understanding of the interaction of Ras with other proteins^10–13^. In fact, the detailed mechanism of Ras-Raf interaction is still a matter of debate.

Ras is a peripheral membrane protein that acts as a switch by binding to GDP or GTP (referred to as Ras.GDP or Ras in this manuscript, respectively). In its GTP bound form, Ras binds to various effector proteins, including Raf^14^. Through a series of yet unclear steps, the binding of Ras to Raf leads to the formation of activated Raf dimers that activates the ERK signaling pathway in eukaryotes, a pathway that controls cell proliferation, cell growth, and cell differentiation, among other important cellular processes. While the role of Raf dimer formation on ERK activation is well-established^15–21^, it is unclear whether Raf-dimer formation is mediated by monomeric Ras or dimeric Ras (Fig 1 A) ^8,22–30^. Recent experiments point in both directions^31–33^. Crucially, the most compelling experiments in favor of Ras dimer formation comes from direct observation using super-resolution microscopy, which has a resolution of 10-20 nm. However, Ras is approximately 2 nm in diameter. Therefore, close association of Ras proteins can be interpreted as Ras dimers or even oligomers even in the absence of any dimers or oligomers (Fig 1 B). Also, the low resolution of the microscope can exaggerate the number of dimers and trimers in cases where Ras truly dimerizes (Fig 1C). Because of which, using direct observation, it is difficult to resolve the debate in favor of one hypothesis or the other.

**Figure 1:**
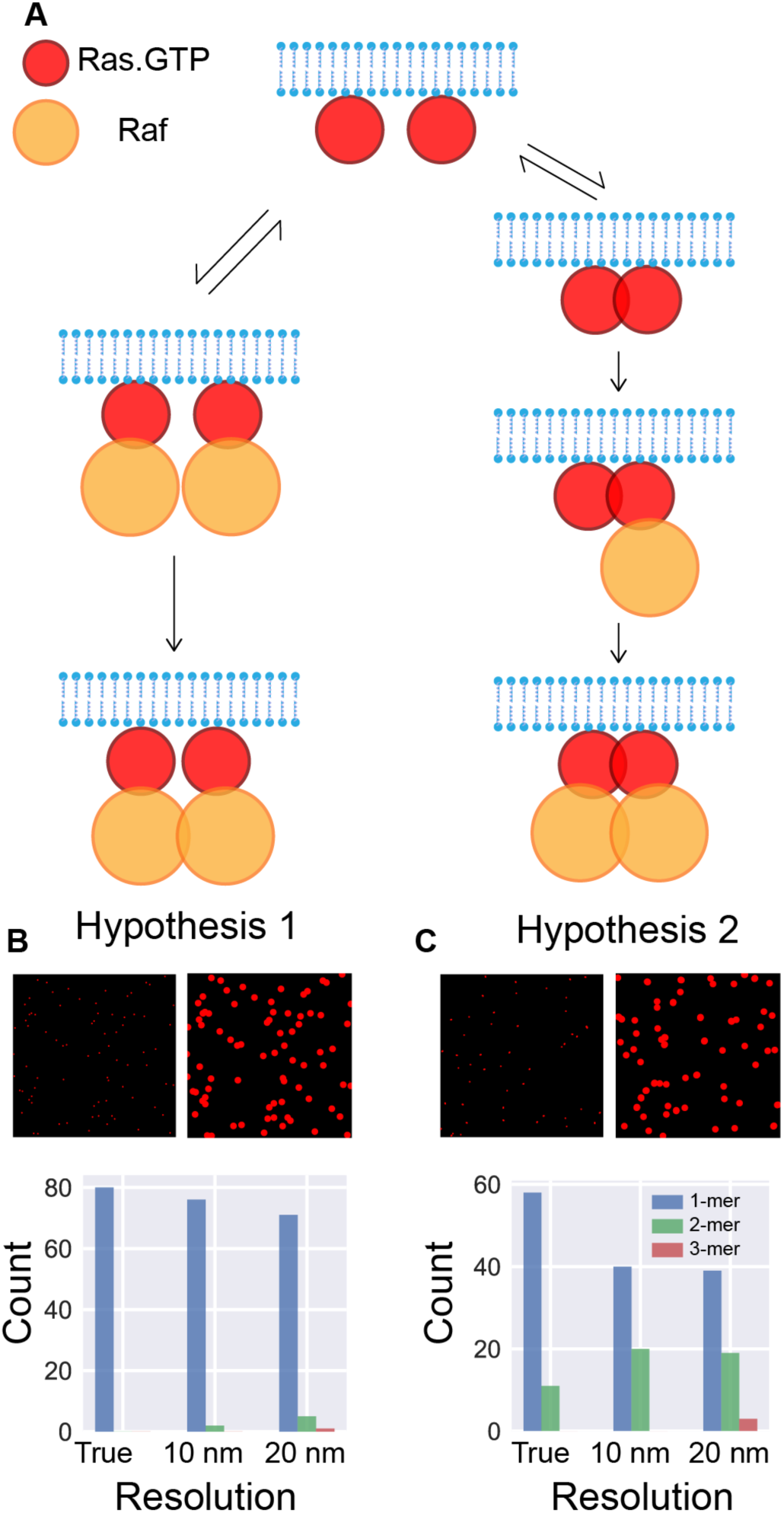
Does Ras dimerize? (**A**) We consider two different hypotheses for Raf activation. In both our hypotheses, Raf activates after forming Ras-Raf hetero-tetramer (last panel). Ras-Raf hetero-tetramer is a Raf dimer, formation of which is necessary for Raf activation^20,21^. The mechanisms for Ras-Raf hetero tetramer formation differ in the two hypotheses. In hypothesis 1, Ras cannot dimerize, but in hypothesis 2, Ras dimerization is essential for the formation of the hetero-tetramer. We also consider a third hypothesis (not shown here) in which both mechanisms are allowed. (**B**) & (**C**) We simulated the dynamics of Ras in the absence of Raf to detect and understand the differences in Ras clustering patterns, if any, generated by the two hypotheses. Surveying the true pattern (top left panel) clearly shows the absence and presence of Ras dimers in hypothesis 1 and 2, respectively. However, when the same point pattern is observed at lower resolution, e.g., 10 nm (top right pattern) or 20 nm, we find spurious Ras-dimers, and even trimers, in the observed point patterns (bottom panel). In light of this observation, because current experimental techniques to detect Ras can, at best, reach 10 nm resolution, we cannot be certain about the presence of Ras-dimers and higher order clusters.

To circumvent this status quo, we propose to resolve this debate using indirect evidences. In this paper, we investigate three hypotheses that investigate Raf dimerization in the presence and absence of Ras dimerization. Experimental evidences show that membrane proteins are subjected to nontrivial diffusive transport^34^. Therefore, assuming that either Ras or Raf is well-mixed may lead to erroneous results and spatiotemporal simulation of their interaction is necessary. We use a recently developed spatiotemporal simulation technique, BD-GFRD^35–41^, to investigate Ras-Raf kinetics in biologically relevant concentrations and timescales. Ras-Raf interaction is endowed with multiple timescales that vary depending on the concentration of Ras and Raf. The concentration dependence of these timescales provides a useful set of probes through which we investigate the indirect effect of the presence or absence of Ras dimerization in Ras-Raf interaction.

## METHODS

### Ras-Raf interaction model

The formation of phosphorylated (activated) Raf dimers is necessary to activate the various downstream effector proteins in ERK pathway. However, the details of the dimerization process are still poorly understood. Experimental data suggests that sometimes Raf dimers form only after forming stable complexes with membrane bound activated (GTP bound) Ras. Unfortunately, most of these measurements are based on optical microscopy, which cannot resolve the interactions between Ras and Raf in the course to form Raf dimers. Because of which, two competing hypotheses has been proposed. In this paper, we investigate the consequences of the underlying assumptions of these hypotheses. In particular, we test how these assumptions influence the activation timescale of Raf dimers and the fraction of activated Raf dimers.

### Hypothesis 1: Ras does not dimerize

The first hypothesis proposes that Ras (*S*) does not form homodimers, and it diffuses freely on the plasma membrane. Raf (*F*) binds to individual Ras and the resultant heterodimer (*SF*) binds with itself to form the Raf dimer (*S*_2_*F*_2_). The Raf dimers phosphorylate each other through colliding with each other^21,42^. The activation kinetics is an assumption of the model. The activation kinetics of Raf remains unclear and currently available experimental data does not rule out activation through interaction of two diffusing *S*_2_*F*_2_s. The following set of chemical reactions represent this hypothesis. For clarity and brevity, we represent Ras and membrane bound Raf through the symbols *S* and *F*, respectively. Also, we represent an activated molecule by adding an asterisk (*) to its symbol.

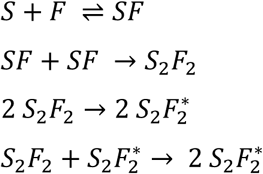

### Hypothesis 2: Ras homodimers are necessary for Raf dimerization

The second hypothesis assumes that free Ras cannot bind to Raf and they need to dimerize before they can bind to Ras. In this hypothesis, Ras forms homodimers (*S*_2_) by binding to another Ras. Two cytosolic Rafs bind to the Ras dimer and form Raf dimers that activate using the same mechanism as in the first hypothesis. The chemical reactions governing second hypothesis are:

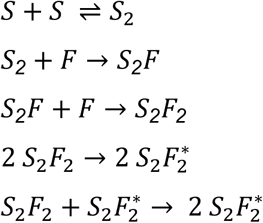

### Hypothesis 3: Both mechanisms present

In this hypothesis we assume that both free Ras and Ras dimers can bind to Rafs. However, a Raf bound Ras cannot form a Ras dimer, i.e., the reaction *SF* + *S* → *S*_2_*F* is not allowed. The activation process is same as the two previous hypotheses. The corresponding chemical reactions are:

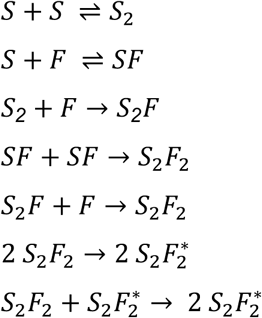

### Simulation Details

#### BD-GFRD

Ras-Raf interactions span multiple timescales: Once Ras and Raf are within interaction distance, formation of Ras-Raf complexes take picoseconds to nanoseconds, but, in biologically relevant concentrations, it usually takes milliseconds or seconds for a Ras and Raf to come within interaction distance. Therefore, any simulation of Ras-Raf interaction should account for both timescales. This requirement rules out all-atom or coarse-grained spatiotemporal simulations, which, at best, can reach milliseconds timescale for a single protein. On the other extreme, well-mixed chemical reaction models can capture all timescales, but leaves out important spatial correlations. Green’s function reaction dynamics (GFRD)^38–40,43^, a recently developed multi-scale method, solves this conundrum. In BD-GFRD^35–37^, an updated version of GFRD, one can perform spatiotemporal simulations that can access seconds and minutes timescale.

BD-GFRD leverages on the fact that biological systems are dilute and biomolecular interactions are short-ranged. These two constraints partition the simulated particles into two groups. Particles that are within interaction radius of each other are simulated using a molecular mechanics algorithm, e.g., Brownian Dynamics (BD). Particles that are far apart from other particles are treated as independent particles and they are propagated diffusively using GFRD, which is an event-driven algorithm. When a particle can be treated as an isolated (non-interacting) entity, GFRD computes the Green’s function for the diffusion equation for that particle with an appropriate boundary condition. We sample the next event time and position from the computed Green’s function. Thus, by simulating the computationally expensive free diffusion using an event-driven algorithm, BD-GFRD can achieve almost a million times speed up compared to all-atom or coarse-grained simulations. *One should note that we can calculate the Green’s functions for spherical particles only*^35,36,43^. Therefore, in our model, we treat every complex as a sphere with radius commensurate with their mass.

The association reactions are modeled as instantaneous processes. When two molecules come within a distance *r*_*asso*_ of each other they react instantaneously and form the protein complex (Fig 2B). We use the same criterion for the activation reactions as well.

**Figure 2:**
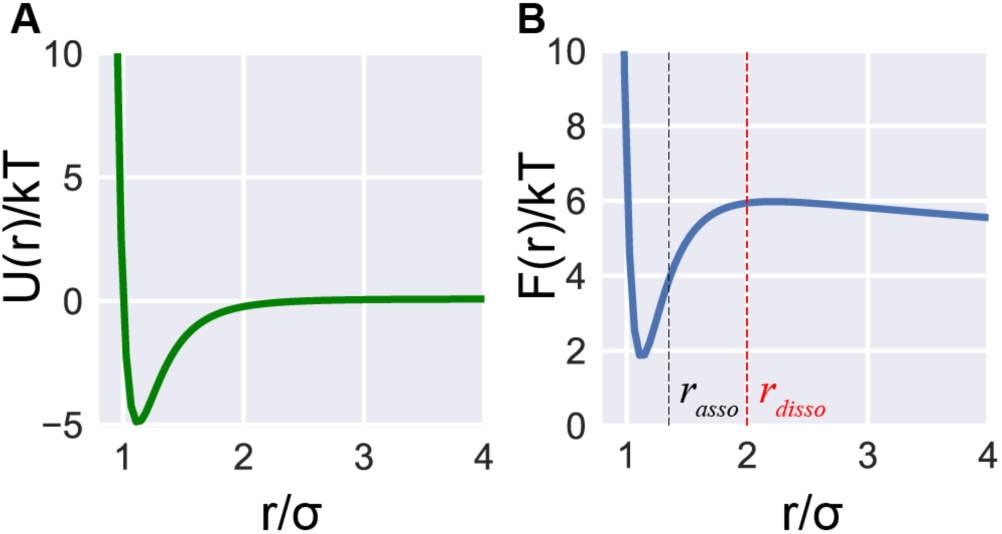
Lennard-Jones Potential. (**A**) Lennard-Jones potential for *ϵ* = 5 *kT*. (**B**) Free energy *F* for the potential in **A**.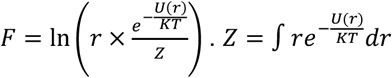 is the partition function in 2D. The free energy is non-monotonic, with a finite barrier at *r* ≈ 2*σ*. Hence, after a dissociation reaction, the dissociated molecules are placed at a distance *r*_*disso*_ = 2*σ* from each other. We chose the minimum distance for association reaction, *r*_*asso*_ ≈ 1.35 *σ*, to ensure that the associating molecules are well below the free-energy barrier.

For dissociation reactions, we assumed that the equilibration of the bond dissociation happens much more quickly than two consecutive dissociation events. Under this assumption, the dissociation events follow a Poisson process and the dissociation times are exponentially distributed. Once two molecules dissociate, they are placed at a distance *r*_*disso*_ from each other. We chose *r*_*disso*_ in such a way to ensure that the dissociated molecules do not immediately bind to each other (Fig 2B).

#### Simulation Parameters

We performed all simulations on a two-dimensional 1 *μm* × 1 *μm* simulation box with periodic boundary condition. All molecules *i* and *j*, except *S*_2_*F*_2_ and 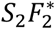, that reacted together in a reaction interacted with each other through isotropic Lennard-Jones interaction with interaction strength, *ϵ*_*ij*_ = 5*kT*, and cutoff radius *r*_*c*_ = 2.5*σ*_*ij*_, where *σ*_*ij*_ = *r*_*i*_ + *r*_*j*_ is the sum of the radius of the two interacting particles. All other molecules, including *S*_2_*F*_2_ and 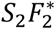, interacted with each other through repulsive WCA potential with *ϵ*_*ij*_ = 3 *kT*. We list the mass and radius of all molecules in the table below.

We used a Langevin integrator at 310 K for the Brownian dynamics. We computed the diffusion constant using Stokes-Einstein relation:

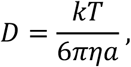

where *η* is the membrane viscosity. We used a membrane viscosity of 120 cP, which is close to the membrane viscosity of eukaryotic cell membranes^44,45^.

We picked *r*_*asso*_ ≈ 1.35*σ*_*ij*_ by examining the free energy landscape of Lennard-Jones potential. Also, we picked *r*_*disso*_ = 2*σ*_*ij*_ using the free energy landscape. We chose the dissociation rate for all dissociation reactions to be 10/s^46,47^.

## RESULTS

### Time evolution of protein concentrations

We start each simulation with a number *n*_*Ras*_ of free Ras (*S*) molecules and a number *n*_*Raf*_ of free Raf (*F*) molecules. Subjected to the reactions in the three hypotheses, *S* and *F* combine to form various intermediate complexes that eventually produce activated *S*_2_*F*_2_ (Figure 3 A-C). Despite the variety of intermediates, a common theme emerges in their time evolution. In all three hypotheses, *S* and *F* concentrations decay to produce the first intermediates that increases in concentrations until the second intermediates form at a time depending on *n*_*Ras*_ and *n*_*Raf*_. In hypothesis 1, for example, *SF* is the first intermediate and *S*_2_*F*_2_ is the second intermediate and in hypothesis 2, *S*_2_ is the first intermediate and *S*_2_*F* is the second intermediate. As the second intermediates form, the concentrations of the first intermediates decrease, peaking at a concentration that also depends on the initial Ras and Raf concentrations. Interestingly, how *S* and *F* decays over time depends on the model hypothesis. Ideally, such an observation would have been useful to identify the nature of Ras-Raf interaction in experiments. However, in biologically relevant concentrations, these variations occur at timescales that are too fast for current experimental probes and too slow for all-atom simulation techniques (Fig 3 A-C).

**Figure 3:**
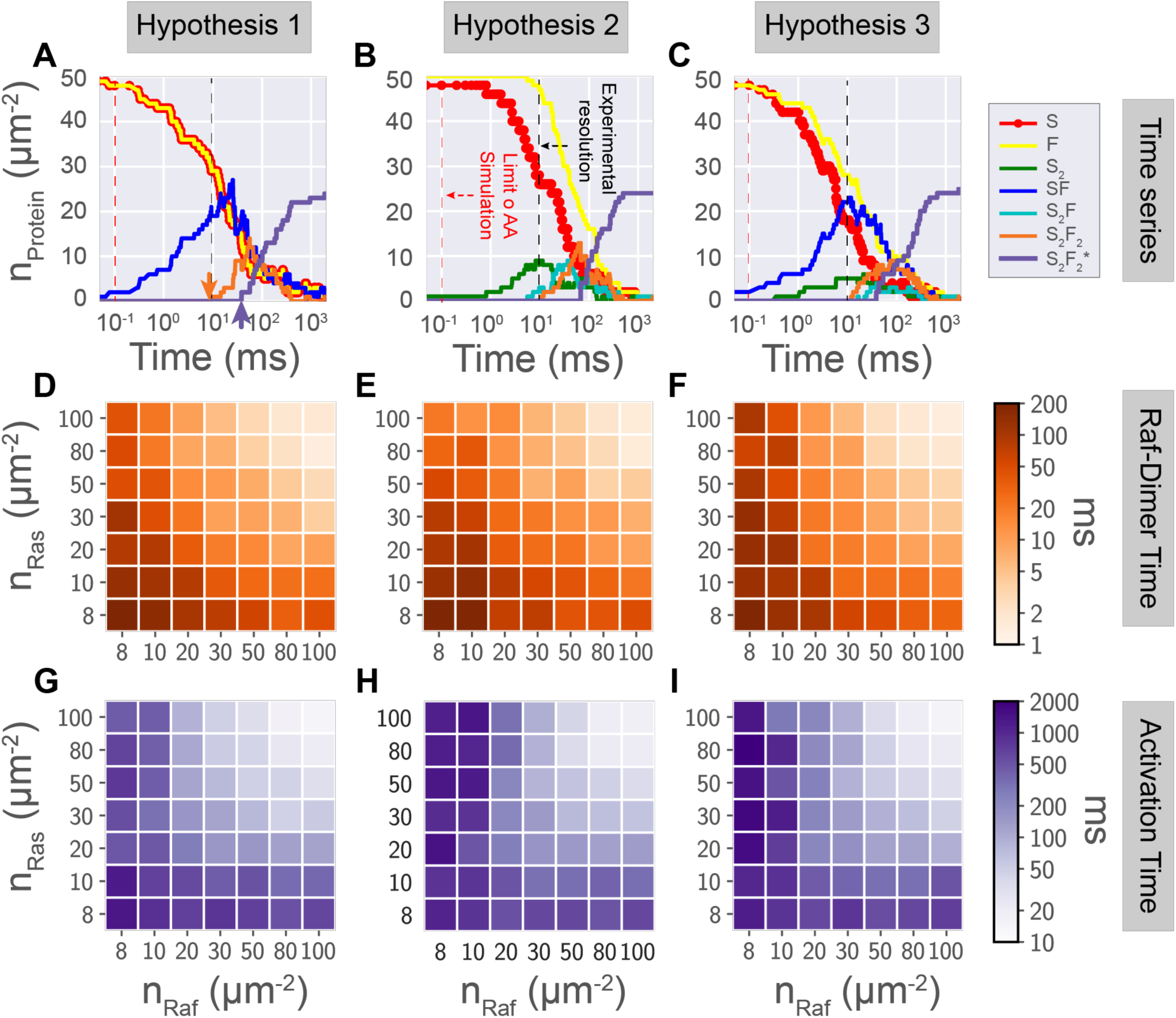
Comparison between hypotheses. (**A**-**C**) Time trace of the concentrations of the reacting molecules. The concentrations, as expected, vary differently under different hypotheses. In particular, the variation of *S* and *F* with time can be a useful probe to differentiate between the three hypotheses. Remarkably, as we had anticipated, there are multiple timescales. Two particularly important timescales are the time at which *S*_2_*F*_2_ first forms, 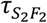 (orange arrow in **A**), and the time at which the first activation of *S*_2_*F*_2_ occurs, *τ*_*A*_ (purple arrow in **A**). For most biologically relevant concentrations, none of these timescales, including 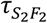 and *τ*_*A*_, can be probed using either experiments or all atom (AA) simulations. We probe how (**D**-**F**) *τ*_*A*_ and (**G**-**I**) 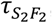 varies with concentrations under different hypotheses. In hypothesis 1 (Ras does not dimerize), the variation is symmetric with respect to *n*_*Ras*_ and *n*_*Raf*_. In contrast, for hypotheses 2 and 3 (Ras dimerizes), this variation is asymmetric. Hence, the (a)symmetry in timescales with variation in concentrations can be used to probe whether Ras dimerizes or not.

Fortunately, Ras-Raf interaction is endowed with multiple timescales that all are potential experimental probes. For example, in hypothesis 1, *S* and *F* combine to produce the Ras-Raf complex, *SF*, which first occurs at a timescale *τ*_*SF*_. *SF* can either irreversibly combine with itself to form the inactive Raf dimer, *S*_2_*F*_2_, or it can reversibly dissociate back to free Ras. Because of which, the concentration of *SF* changes non-monotonically with time and peaks at a time, *τ*_*peak*_, which depends on *n*_*Ras*_ and *n*_*Raf*_. Also, because of the competition between the dissociation and the combination reactions, *S*_2_*F*_2_ forms at a timescale 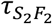 and the first activation of Raf dimers happen at a timescale *τ*_*A*_. Understandably these timescales depend on *n*_*Ras*_ and *n*_*Raf*_ Characterizing the concentration dependence of these timescales is important to construct a dynamic picture of Ras-Raf interaction and to develop them as potential experimental probes for Ras-Raf interaction.

Two different timescales, 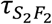 and *τ*_*A*_, are common to all three hypotheses. Hence, we compare and contrast them here. The Raf-dimer formation varies across the three hypotheses, whereas, the activation process remains same. Therefore, using these two timescales, we can probe respectively the direct and the indirect consequences of our assumptions in the three hypotheses. To understand these relationships, we probe these two timescales over different Ras and Raf concentrations. Visually, these two timescales behave differently under different hypothesis (Figure 3 DF & G-I).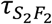 varies symmetrically with *n*_*Ras*_ and *n*_*Raf*_ for both hypotheses 1 and 2, but *τ*_*A*_ varies symmetrically only for hypothesis 1. The variation of both timescales is asymmetric in other cases. Particularly, *τ*_*A*_ varies asymmetrically for hypotheses 2 and 3, both of which admit Ras dimerization. In the absence of Ras dimerization, *τ*_*A*_ varies symmetrically. *Therefore, it is tempting to declare the presence and absence of asymmetry in τ*_*A*_ *as an indirect probe for the presence and absence of Ras dimers. To embolden this claim we investigate the kinetics of the Raf-dimer formation reaction and the activation reaction.*

### Concentration dependence of timescales

To disentangle how *n*_*Ras*_ and *n*_*Raf*_ influence the timescales, we study their individual contribution on the timescales 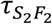 and *τ*_*A*_. We vary either *S* or F concentrations, keeping the concentration of the other molecule fixed. We refer to the former as the **probe** and the later as the **control** molecule.

#### Variation of 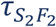

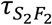 varies similarly with *n*_*Ras*_ (Fig 4 A-C) and *n*_*Raf*_ (Fig 4 D-F) across the three hypotheses. In all three cases, 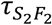 decreases with increasing probe molecule concentration, eventually saturating to a timescale dependent on the control molecule concentration. For example, for *n*_*Ras*_ = 8 and in hypothesis 1, 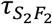 decays from a maximum value of ∼240 ms for *n*_*Ras*_ = 8 to ∼ 50 ms for *n*_*Raf*_ = 50 and stays there till *n*_*Raf*_ = 100, the highest concentration probed (Fig 4A). Although the data for Fig 4F is noisy, the decreasing trend is easily discernible. The decay rate of 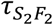 with the probe concentration is same across different control molecule concentration, and across different hypothesis. In contrast, the asymptotic timescale, i.e. the timescale at large probe molecule concentration, varies similarly with both control molecule concentration under a hypothesis, but the variation is different across hypothesis. For example, in Fig 4G, the asymptotic 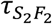 varies as *n*^−1.5^, independent of the chosen control molecule; *n* is the concentration of the control molecule. However, asymptotic 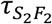 varies as *n*^−1.3^ and *n*^−1.2^ for hypotheses 2 and 3, respectively. Thus, the effect of the hypothesis is directly reflected in the variation of asymptotic 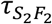 with respect to the concentration of the control molecule. It is unclear what processes determine the observed scaling. We suspect the underlying reaction kinetics is the main determinant of the timescales, because if these timescales resulted from a diffusion limited process, then the timescale would decay as *n*^−1^, which is the well-known Berg-Purcell scaling^48^.

**Figure 4:**
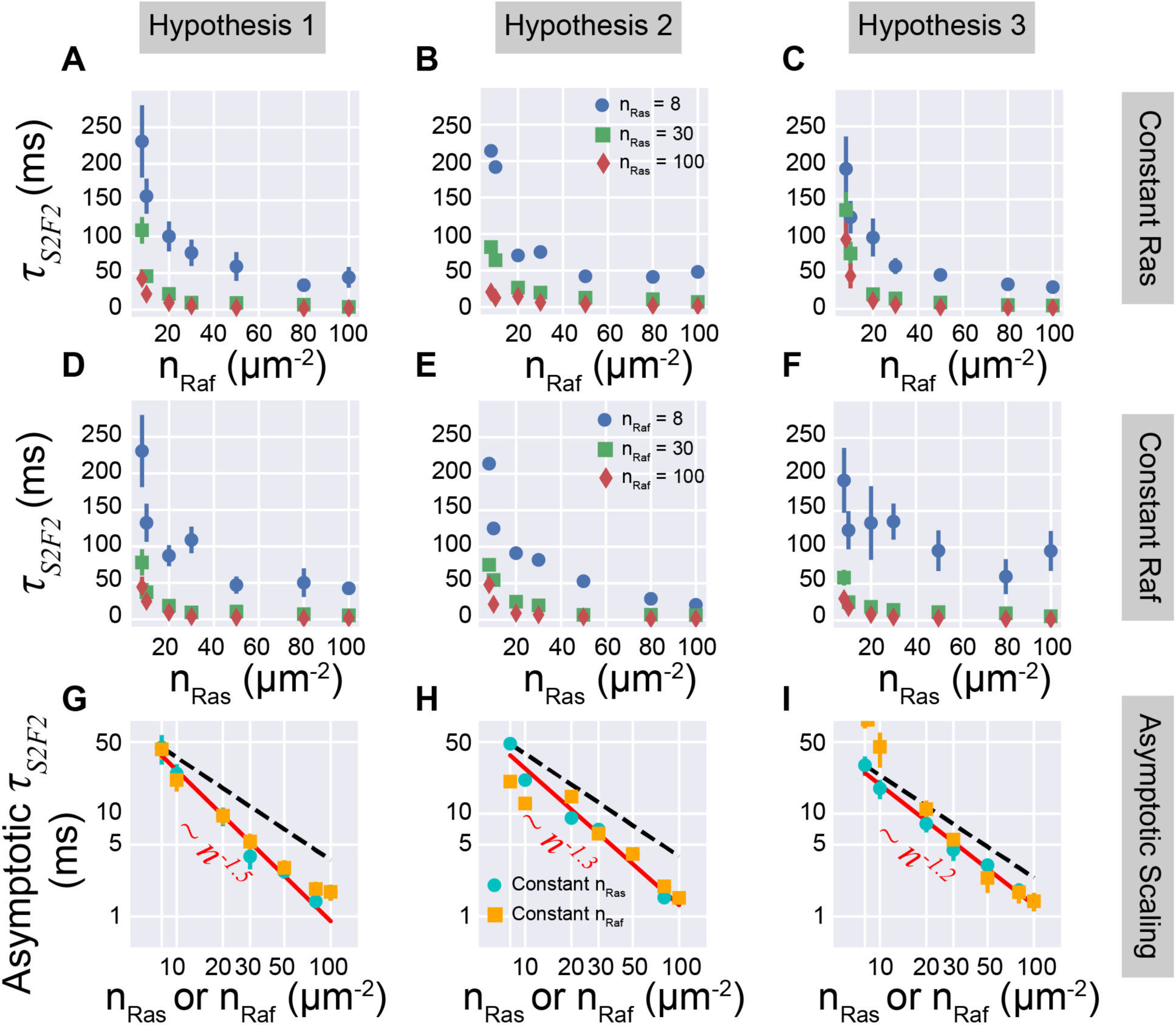
Concentration dependence of 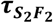. We probed Raf dimer formation timescale, 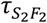, (**A**-**C**) by varying *n*_*Raf*_, keeping *n*_*Ras*_ constant and (**D**-**F**) by varying *n*_*Ras*_, keeping *n*_*Raf*_ constant. For both cases, 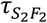 varied uniformly across the three hypotheses: 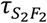 decreased with increasing *n*_*Raf*_ and *n*_*Ras*_, eventually saturating to an asymptotic timescale that depends on the protein concentration that is kept constant (*n*_*Ras*_ for (**A**-**C**) and vice versa). Although the data is noisy for (**F**) a decreasing trend is not difficult to infer. (**G**-**I**) The asymptotic 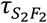 is not diffusion limited, because the timescale does not decrease linearly with concentration (black dashed line). Instead, the asymptotic timescale decreases as a power law 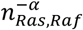 (red line), where *α* depends on the hypothesis (see figure). What determines the value of *α* is unclear, but we hypothesize that the difference arises from the difference in the underlying kinetics.

**Figure 5:**
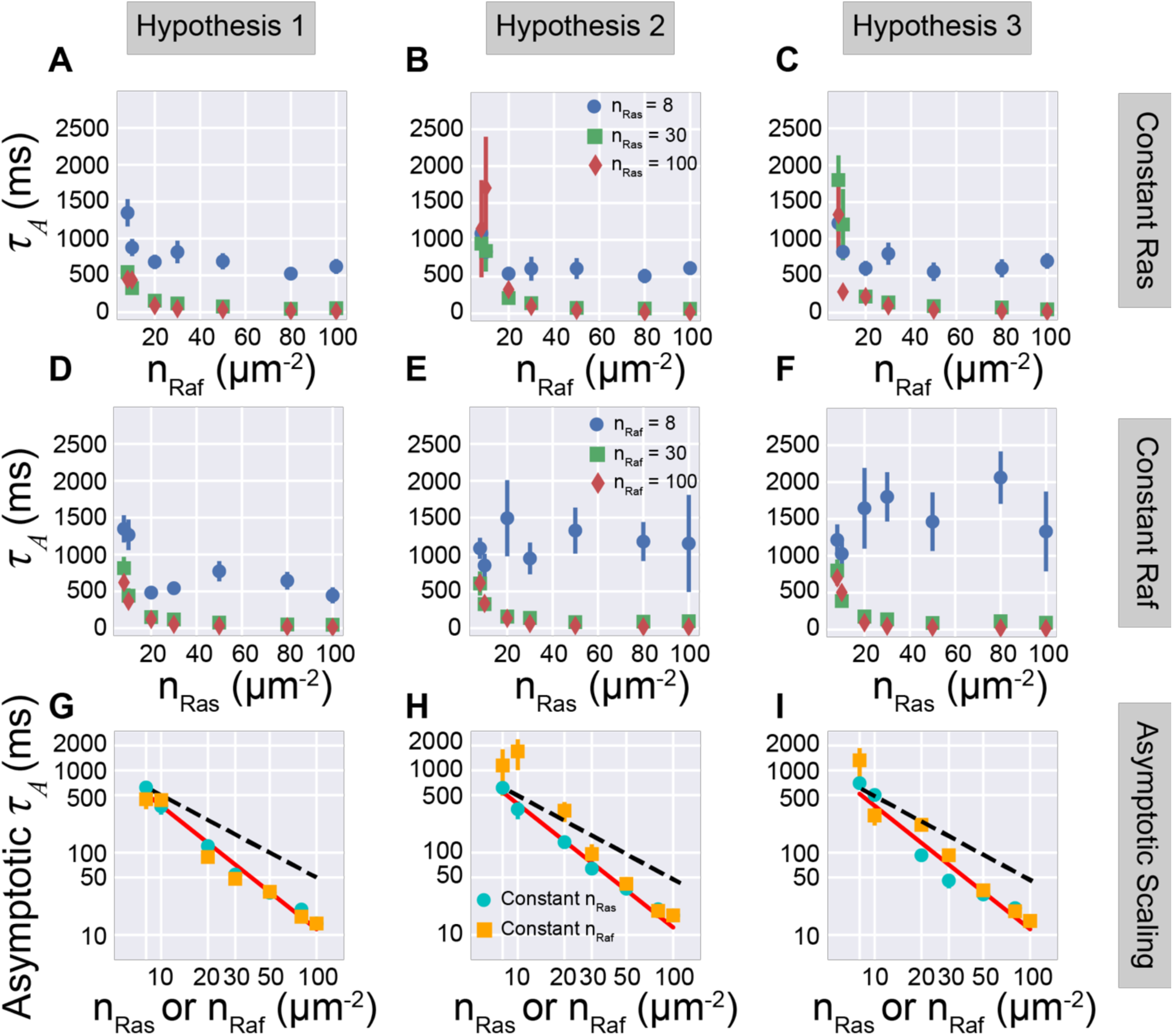
Concentration dependence of activation timescale. We probed activation timescale (**A**-**C**) by varying *n*_*Raf*_, keeping *n*_*Ras*_ constant and (**D**-**F**) by varying *n*_*Ras*_, keeping *n*_*Raf*_ constant. For (**A**-**C**), the activation timescale varied uniformly across the three hypotheses: *τ*_*A*_ decreased with increasing *n*_*Raf*_, eventually saturating to an asymptotic timescale that depends on *n*_*Ras*_. In contrast, for (**D**-**F**), *τ*_*A*_ varied differently across the three hypotheses. For hypothesis 1, *τ*_*A*_ decreased with increased concentrations for all values of *n*_*Raf*_, but *for hypotheses 2 and 3 τ*_*A*_ *increased with increasing n*_*Ras*_ *at low values of n*_*Raf*_. Although, at higher values of *n*_*Raf*_, *τ*_*A*_ behaved similarly to hypothesis 1. (**G**-**I**) The asymptotic *τ*_*A*_ is not diffusion limited, because the timescale does not decrease linearly with concentration (black dashed line). Instead, the asymptotic timescale decreases as 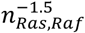 (red line) for all three hypotheses, implying that the activation time is set by a common kinetic process.

#### Variation of *τ*_*A*_

Unlike 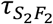, *τ*_*A*_ varies differently with *n*_*Ras*_ and *n*_*Raf*_ across the three hypotheses. For hypothesis 1, *τ*_*A*_ decays with both probe concentrations, and asymptotes to a value depending on the control concentrations, in a fashion similar to 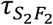. In contrast, for hypotheses 2 and 3, the variation with respect to probe molecules depend on the choice of the probe. If the probe molecule is *F*, and the control molecule is *S*, then *τ*_*A*_ decays with increasing concentration and asymptotes to a value that only depends on *n*_*Ras*_ However, if *S* is chosen as the probe and *F* as the control, then for low *n*_*Raf*_ *τ*_*A*_ *increases* with increasing *n*_*Ras*_ and asymptotes to a value depending on *n*_*Raf*_. Remarkably, in both hypotheses 2 and 3, Ras dimerizes, whereas in hypothesis 1, Ras does not dimerize. Does Ras dimerization cause the observed anomalous variation of *τ*_*A*_?

To answer this question, we investigated how the time series of concentrations for hypothesis 2 and 3 at low concentrations of *F* (Fig 6). We find that for hypothesis 2, the delay results from the formation of *S*_2_*F*, which forms from the *irreversible* association of a Ras dimer, *S*_2_, and free Raf, *F*. When *n*_*Ras*_ is low, *S*_2_*F* forms with less propensity and enough *F* remains available to form at least two *S*_2_*F*_2_, the minimum number of inactive Raf dimer required for the activation reaction. As *n*_*Ras*_ increases, *S*_2_*F* forms with increasing propensity, resulting in decreasing number of *F* available in the simulation box. Because of which, the propensity of the formation of *S*_2_*F*_2_ decreases with increasing *n*_*Ras*_ and the activation timescale increases. A slightly different mechanism causes the anomalous variation in hypothesis 3. In this hypothesis, as *n*_*Ras*_ is increased, the propensity of *SF* and *S*_2_*F* formation increases. Because of which, free *F* remains present in low numbers until *SF* dissociates back into free *S* and *F*. A lack of free *F* decreases the propensity of the formation of *S*_2_*F*_2_ that increases the activation time (Fig 6 and S1). Therefore, resource limitation caused by the low concentration of *F* is the principle reason behind the anomalous behavior. However, the low concentration of *F* affects the activation timescale because Ras-dimerization mediated two-step Raf-dimer formation mechanism (*S*_2_+ *F* → *S*_2_*F, S*_2_*F* + *F* → *S*_2_*F*_2_) requires more freely available *F*. Even if the formation of *S*_2_*F* were reversible in both hypotheses, we would have similar delay, because, in such a case, we would have to wait until the release of *F* from *S*_2_*F. Hence Ras-dimerization is the underlying cause of the observed anomalous behavior.*

**Figure 6:**
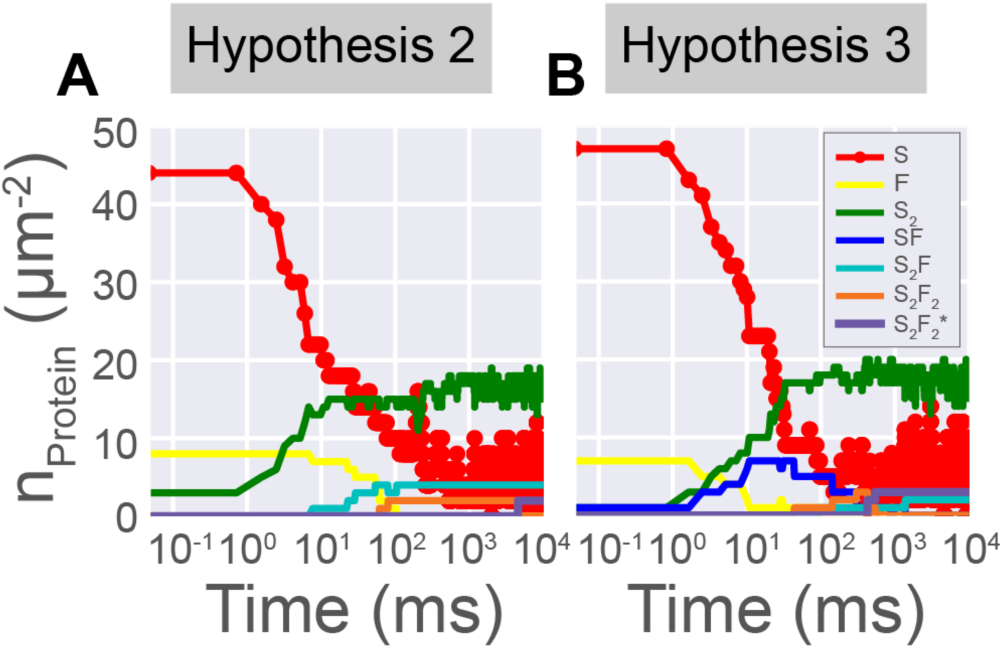
Activation events in hypotheses 2 and 3. For low concentrations of Raf (8/*μm*^2^here), in hypotheses 2 and 3 activation time increases with increasing Ras concentration. The underlying reason for this anomaly is the limited availability of *F*, which happens due to different reasons in the two hypotheses. (**A**) In hypothesis 2, *F* becomes limited due to the formation of *S*_2_*F*, a molecule necessary to form *S*_2_*F*_2_. (**B**) In hypothesis 3, *F* readily forms *SF*. However, due to the higher concentration of *S*, the preferred route to form *S*_2_*F*_2_ is through the formation of *S*_2_*F*, which can occur only after *SF* dissociates back to *S* and *F*. The higher the concentration of *S*, the higher the propensity to form *SF*. Because of which, activation time increases with increasing Ras concentration.

Although the variation of *τ*_*A*_ with respect to probe concentration depends on the underlying hypothesis, the variation of asymptotic activation time with control molecule concentration remains uniform across the three hypotheses! In all three cases, the asymptotic timescale varies as *n*^−1.5^, where, like 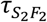, *n* is the concentration of the chosen control molecule. Again, the asymptotic timescale does not follow Berg-Purcell scaling, implying that the underlying reaction kinetics determine the asymptotic timescale.

### Reaction kinetics

Although the molecules in our simulation box move diffusively, as we have seen, their reaction timescales differ significantly from the limit set by pure diffusion. Hence, we suspect that the underlying reaction kinetics set the observed timescale. Because the activation timescale varies uniformly across the three hypotheses, we investigate its variation with the control molecule concentration, *n*.

In our models, a *S*_2_*F*_2_ molecule gets activated when it collides with another *S*_2_*F*_2_ or 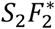 molecule. However, because the activation time is the time at which the first activation reaction happens, its kinetics is entirely determined by the concentration of *S*_2_*F*_2_, [*S*_2_*F*_2_]. In particular, we find that the concentration of *S*_2_*F*_2_ right before the activation event is related to the inverse of the activation time through a simple well-mixed mass-action like form. That is the activation rate, *r*_*A*_ = 1/*τ*_*A*_ is proportional to the mass-action flux, 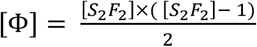:

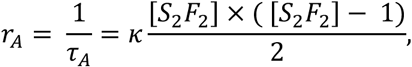

where *κ* is the rate constant. This relationship holds uniformly across the three hypotheses, as we show in Fig 7A. In fact, the rate constant *κ* is exactly equal to the inverse of the activation time if there were only two *S*_2_*F*_2_ (≈ 6 × 10 ^−4^ *ms*^−1^*μm*^4^). The uniformity of the activation kinetics and the uniformity of the power law exponent of the asymptotic *τ*_*A*_ across three hypotheses lends credence to our suspicion that the underlying reaction kinetics sets the timescale. If this suspicion is true, then the kinetics of *S*_2_*F*_2_ should be different across the three hypotheses. We find this to be the case for hypotheses 1 and 2 (Fig 7B). We could not compare hypothesis 3 with the other hypotheses because *S*_2_*F*_2_ forms through two different processes in hypothesis 3. Therefore, the total mass action flux is a weighted sum of the two fluxes: [Φ] = *κ*_l_[Φ_l_] + *κ*_2_[Φ_2_]. Hence, the rate constant cannot be compared in a straightforward manner, like the other two hypotheses.

**Figure 7:**
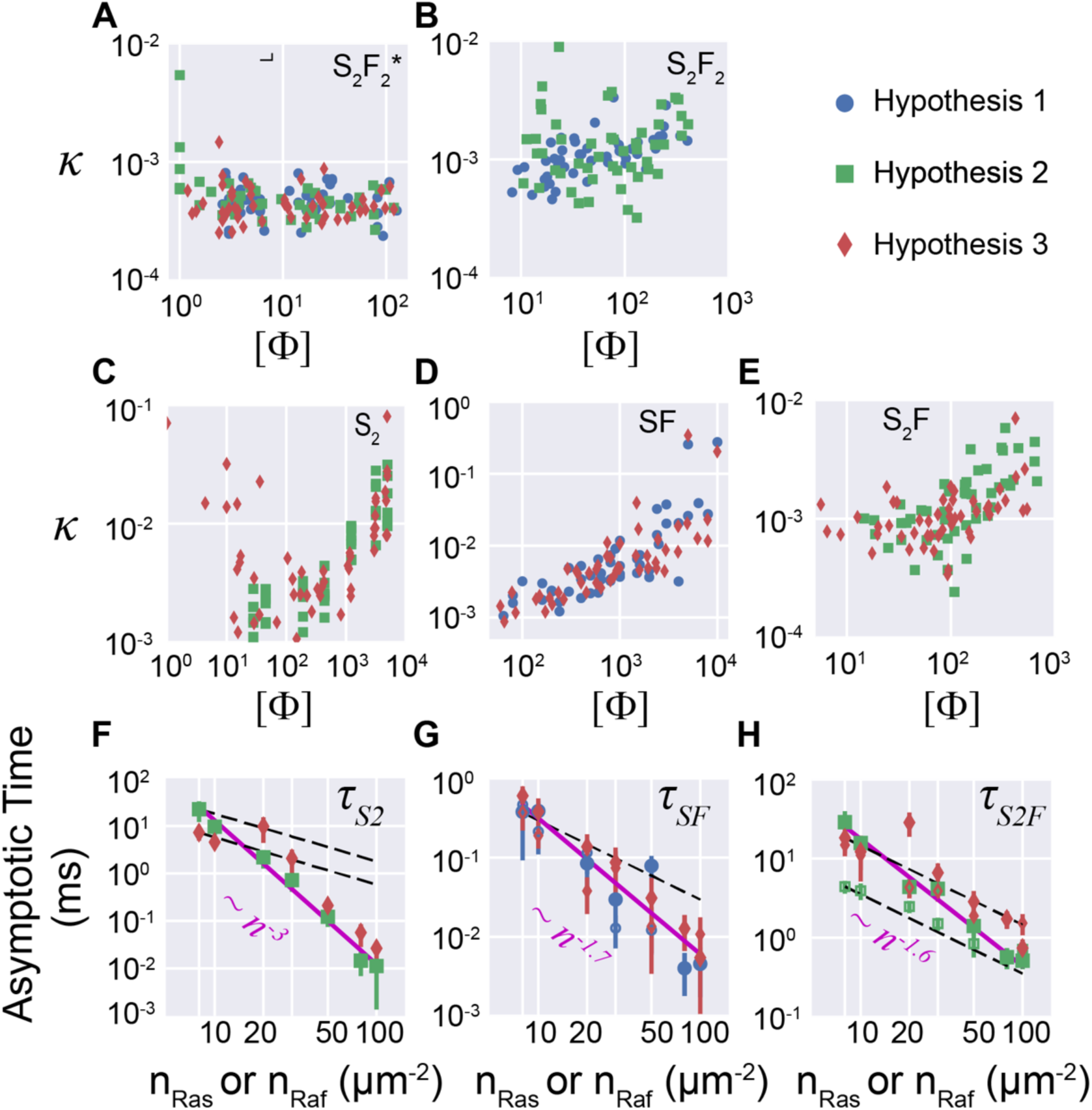
Comparison with mass-action kinetics. (**A**) Independent of the hypothesis, the mean activation rate (1/*τ*_*A*_) follows mass action kinetics, 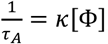, for a wide range of concentrations as shown by the insensitivity of the reaction rate constant *κ* = 1/(*τ*_*A*_ × [Φ]), on the mass action flux 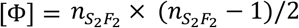. (**B**-**E**) In contrast, other timescales do not arise from well-mixed mass action kinetics, because *κ* depends on the mass action flux [Φ]. Mass action flux for various reactions are described in the text. (**F**-**H**) The deviation from the mass-action behavior is reflected in the asymptotic scaling of the timescales. For example, the difference in the deviation from mass-action behavior in **C** is reflected in the different asymptotic behavior in **F**. In contrast, the similarity of deviation from mass-action kinetics in **D** is reflected in the same asymptotic behavior in **G**. Solid markers for variations with respect to *n*_*Ras*_ and open markers for *n*_*Raf*_. For *SF*, there is no difference in variation with respect to *n*_*Ras*_ or *n*_*Raf*_. *S*_2_*F*, however, shows different variation. Most interestingly, the asymptotic timescale follows Berg-Purcell scaling for all the cases except variation with respect to *n*_*Ras*_ in hypothesis 2.

We consolidated the connection between the asymptotic scaling and underlying kinetics by repeating the above analysis for other molecules (Fig 7 C-H). We list the mass action fluxes for the corresponding reactions in Table 2. We found that the asymptotic scaling of the timescales reflects the deviation from mass-action kinetics. For example, in Fig 7C, the deviation from mass-action like behavior is different for hypothesis 2 and 3. For hypothesis 2, *κ* increases monotonically with [Φ]. In contrast, for hypothesis 3, *κ* decreases with [Φ] initially and then it increases with [Φ] with the same functional form as in hypothesis 2. This difference is also reflected in the scaling of asymptotic 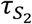 with *n*_*Ras*_. For hypothesis 2, we observe a single power law. However, for hypothesis 3, the scaling is Berg-Purcell like in the beginning, which changes to the same power law we observe for hypothesis 2. We observe a similar correlation between the underlying kinetics and the scaling of the asymptotic timescale for *SF* and *S*_2_*F*.

**Table 1:** Mass of the protein complexes involved in Ras-Raf interaction and the radius of the spheres representing the complexes. We assumed that the densities of the proteins are 1 gm/cc. We approximated the diffusion constant using Stokes-Einstein relationship: 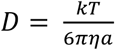, by assuming that the protein is a sphere of radius *a* and is embedded in an isotropic fluid of viscosity *η* = 120 cP.

| Molecule | Mass (kDa) | Radius of a sphere of equal mass, $a$ (nm) | Diffusion coefficient ($\mu m^2/s$) |
| --- | --- | --- | --- |
| $S$ | 22 | 2.0 | 0.95 |
| $F$ | 66.5 | 2.9 | 0.66 |
| $S_2$ | 44 | 2.5 | 0.76 |
| $SF$ | 88.5 | 3.16 | 0.60 |
| $S_2F$ | 110.5 | 3.43 | 0.55 |
| $S_2F_2$ and $S_2F_2^*$ | 177 | 4.0 | 0.48 |

**Table 2:** Mass-action flux: Flux of the reactants to produce the molecule listed on the left-most column, as modeled by the well-mixed mass action kinetics. Some reactions are not available in all three hypotheses. Where they are absent, we mark the flux by “-” sign. † *S*_2_*F*_2_ forms through two different mechanisms. Hence, the mass-action flux does not have a simple form. So, we did not compute the mass-action flux for hypothesis 3.

| Molecule | | $[\Phi]$ | |
| --- | --- | --- | --- |
|  | Hypothesis 1 | Hypothesis 2 | Hypothesis 3 |
| $SF$ | $[S][F]$ | - | $[S][F]$ |
| $S_2$ | - | $[S] \times ([S] - 1)/2$ | $[S] \times ([S] - 1)/2$ |
| $S_2F$ | - | $[S_2][F]$ | $[S_2][F]$ |
| $S_2F_2$ | $[SF] \times ([SF] - 1)/2$ | $[S_2F][F]$ | † |
| $S_2F_2^*$ | $\frac{[S_2F_2] \times ([S_2F_2] - 1)}{2}$ | $\frac{[S_2F_2] \times ([S_2F_2] - 1)}{2}$ | $\frac{[S_2F_2] \times ([S_2F_2] - 1)}{2}$ |

## DISCUSSION

Using a novel spatiotemporal simulation technique, we have investigated how Ras dimerization affects Ras-Raf interaction. We have compared and contrasted the kinetics of three hypotheses that consider Ras-Raf interaction kinetics in the absence or presence of Ras dimerization. As expected, we find that multiple timescales are a feature common to all three hypotheses. These timescales depend on the initial concentrations of Ras and Raf and the presence or absence of Ras dimers affect this trend. We find that when Ras does not dimerize (hypothesis 1), the activation timescale decreases with increasing concentrations of both Ras and Raf. If Ras dimerizes, then this symmetry in Ras and Raf concentrations no longer hold. Instead, if Ras concentration is increased keeping Raf concentration low and constant, then the activation timescale *increases with increasing Ras concentration*. The underlying reason for this reversal in the trend is resource limitation. Free Raf, *F*, takes part in two different reactions when Ras dimerizes. When the concentration of *F* is low and concentration of *S* is high, all *F* are used to form the intermediate molecules, such as *SF* or *S*_2_*F*, and no *F* is left to produce *S*_2_*F*_2_ from these intermediates. *S*_2_*F*_2_ can form if only *F* becomes available again through the dissociation of some of these intermediate molecules. *Although these results are dependent on the particular details of our hypothesis, we find this opposite trend in the concentration dependence of the activation time whenever Ras dimerizes. Therefore, we may use the concentration dependence of the activation time as an indirect probe for Ras dimerization.* These predictions can be tested in *in vitro* experiments^33^, where the concentrations of Ras and Raf can be independently controlled. Due to the simplicity of our system, these predictions may not be directly tested *in vivo*, where unusually low concentration of Ras may render the cell-line nonviable.

The kinetics of the Raf dimer activation follow well-mixed mass action kinetics. We find this observation to be the *exception*, rather than the norm because the kinetics of the formation of the intermediate complexes *do not obey mass-action kinetics*. The underlying reason for such strange reaction kinetics is unclear, but we suspect two probable factors combine to produce such unusual kinetic behavior: (a) inter-particle interaction and (b) reaction timescales. Firstly, every other reaction except the activation reaction is an association reaction, where the reacting molecules interact with each other through attractive Lennard-Jones potential. In the activation reaction, the inactive Raf dimers interact through repulsive WCA interaction, which can be approximated as hard-sphere interaction. It has been suggested that mass-action kinetics works only when the interacting particles interact through hard-sphere interaction^49^. In the presence of attractive interaction, mass action kinetics breaks down. Secondly, the activation reactions occur at a timescale long enough to explore the simulation box completely. Hence, the well-mixed assumption is justified. In contrast, other reactions occur at timescales that are orders of magnitude smaller than the activation timescale. It is impossible to explore the entire simulation box in such a short time and the well-mixed approximation breaks down. It will be interesting to disentangle the role of the timescale and the interaction in the Ras-Raf kinetics. In particular, it will be useful to understand under what conditions our results are comparable to a well-mixed reactor where reactants interact with mass-action kinetics.

Despite the simplicity of our model, it has nontrivial kinetic behavior which are both biologically and chemically interesting. It is likely that our hypotheses will be limited in scope to *in vitro* studies, where Ras-Raf interaction can be studied in isolation. In the presence of other interacting proteins, for example *in vivo* studies, the kinetics is likely to be far richer than what we have observed here. In particular, the competition between Raf, PI3K, PLCε and other effectors of Ras will lead to interesting resource limited kinetics spanning multiple timescales, potentially similar to what we have observed here. The main challenge lies in studying these proteins in biologically relevant conditions. Proteins interact with other proteins through highly anisotropic and specific forces, which we have ignored in our current model by choosing isotropic interactions of identical strength. The presence of anisotropy and specificity can vastly change the timescales of interaction and the resultant kinetics. Also, proteins can indirectly interact with other proteins through the lipids present in the membrane. Simulating the lipid dynamics in conjunction with protein dynamics is beyond the scope of BD-GFRD. Hence, we have to incorporate the effect of lipids through some effective contribution. For example, in this work, we have incorporated their effect through the viscosity of the membrane. We envision to incorporate the anisotropy and the specificity of protein interactions in our model. However, the intended purpose of BD-GFRD is not to simulate biomolecular interactions with detailed biomolecular interactions. Specialized simulations^50^ are in development to answer these questions, which can accommodate detailed biomolecular interactions but fail to reach experimentally accessible timescales. BD-GFRD takes a complementary approach by trading biological details for long timescale simulation. In this way, these very different approaches can complement each other’s findings. For this work, we found it necessary to keep our model simple to understand the role of Ras dimerization in Ras-Raf interaction. Despite these simplifications, the predictions from our model are quite general and experimentally testable with available technologies.

To conclude, Ras dimerization affects the kinetics of Ras-Raf interaction and this influence is reflected in the long timescale behavior of the underlying molecular concentrations. Our results offer a well-defined and well-characterized null hypothesis through which Ras-Raf interaction can be probed to its full extent.

## Acknowledgement

This work was supported in part by Los Alamos National Laboratory LDRD funds through the Center for Nonlinear Studies. This work has also been supported in part by the JDACS4C program established by the U.S. Department of Energy (DOE) and the National Cancer Institute of the National Institutes of Health. Computations used resources provided by the LANL Institutional Computing Program, which is supported by the U.S. DOE National Nuclear Security Administration under Contract No. DE-AC52-06NA25396. The authors acknowledge discussions with T. Bhattacharya, C. Neale, A. Voter, W. Hlavacek, Y.T. Lin, and S. Gnanakaran at LANL, L. Sbailo (Berlin), and D. Goswami (NCI).

## Supplementary Information

**Fig S1:**
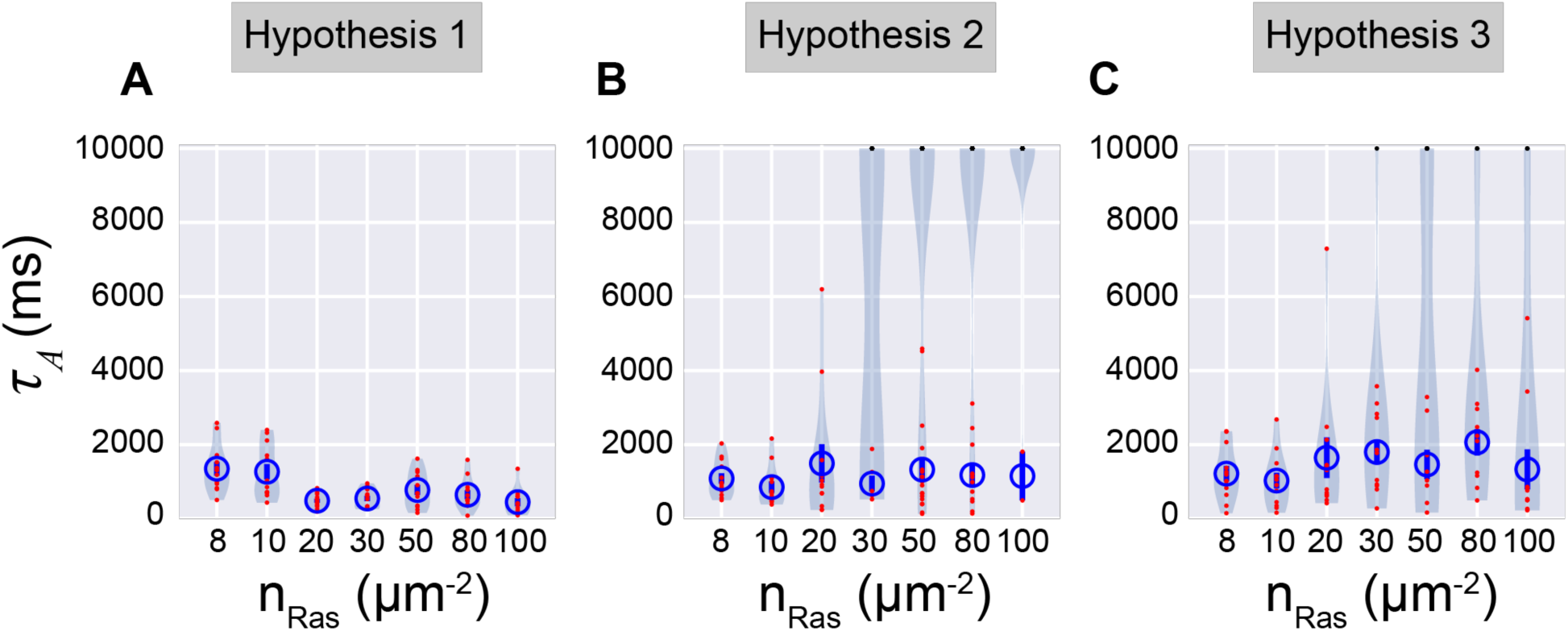
Distribution of activation time for *n*_*Raf*_ = 8. The distribution of the activation time for constant *n*_*Raf*_ for the three hypotheses is displayed here using violin plots. The scatter points along the violin plot shows the raw data. (A) For hypothesis 1, the distribution of the activation time is narrow for all values of *n*_*Ras*_ and the mean value of the activation time (open blue circles) decreases with increasing *n*_*Ras*_ However, for hypotheses 2 (B) and 3 (B), at higher *n*_*Ras*_, the distribution of *τ*_*A*_ is broad. In fact, for a significant fraction of runs, we did not observe any activation events within the time limit of the simulation (10s). We have represented these scatter points with black dots and have not used them to calculate the mean activation time. In both hypotheses 2 and 3, the mean activation time does not decrease with increasing *n*_*Ras*_. In fact, for hypothesis 3, it clearly increases with *n*_*Ras*_.

